# Regulatory T cell stability determines the efficiency of bile duct regeneration during cholangitis

**DOI:** 10.1101/2025.04.02.646748

**Authors:** Naruhiro Kimura, Man Chun Wong, Gareth Hardisty, Ben Higgins, Pei-Chi Huang, Maxym Besh, Daniel A Patten, Timothy Kendall, Prakash Ramachandran, Adriano G Rossi, Graham Anderson, Shishir Shetty, David Withers, Atsunori Tsuchiya, Shuji Terai, Wei-Yu Lu

## Abstract

**Background and Aims:** Cholangiopathies such as Primary Sclerosing Cholangitis (PSC) cause damage to the bile ducts and fibrosis with no effective cure. A reduction and impairment in the function of regulatory T cells (Tregs) occurs in PSC. Yet, it is currently unknown what consequence this has on bile duct regeneration. We investigate whether Tregs regulate bile duct regeneration and the dynamics of Tregs turnover during bile duct injury.

**Approach and Results:** We used the transgenic Foxp3^GFPDTR^ model to mimic reduced Tregs infiltration to the liver during bile duct injury and showed that reduced intrahepatic Tregs limits bile duct regeneration. Fate mapping of Tregs (Foxp3^CreERT^Ai14) showed that Tregs acquire a pro-inflammatory phenotype within the inflammatory microenvironment, even after IL2 mediated Tregs expansion. Ox40L expression correlates with fibrosis in PSC patients. In the 3,5-Diethoxycarbonyl-1,4-Dihydrocollidine (DDC)-diet mouse model of experimental cholangiopathy, combining IL2-complex administration and blocking Ox40L decreases periportal fibrosis level, increases Treg number and reduces the pro-inflammatory phenotype of Tregs.

**Conclusions:** These results indicate that Tregs mediate cholangiocyte response to biliary injury, and Tregs downregulate Foxp3 and acquire an inflammatory phenotype in an inflammatory microenvironment. Enhancing Treg numbers through IL2 complex administration and blocking Ox40 signalling simultaneously suppress the inflammatory phenotype of Tregs and reduces bile duct damage and fibrosis.

## Introduction

In severe and chronic liver diseases including cholangiopathies such as Primary Biliary Cholangitis (PBC) and Primary Sclerosing Cholangitis (PSC), the development of a ductular reaction (DR) within the liver is one of the compensatory liver repair mechanisms, with hepatic progenitor cells (HPC) of biliary origin that reside within the bile ductules being activated during DR^1–3^. Interestingly, the magnitude and phenotype of DR in human liver regeneration are variable depending on disease pathogenesis and the composition of the regenerative niche. For example, DR observed in PBC patients is more robust and proliferative compared to PSC patients despite both diseases being categorized as cholangiopathies ^4^. Besides the regenerative role of DR, the emergence of DR also links with tissue scarring and fibrosis. The dynamics of DR are affected by the tissue microenvironment, especially the immune compartment comprising of macrophages, neutrophils and lymphocytes^5^.

The exact cause of PSC is unknown, although recent genome-wide associated studies (GWAS) and clinical studies have indicated that PSC is immune-mediated ^6^ ^7^, with risk loci identified involved in pathways of T cell regulation such as IL2 and TRAF signalling ^8^. Furthermore, regulatory T cells (Tregs) that suppress immune responses are reduced in PSC patients^9^, which may be one factor contributing to the pathogenesis of PSC. Although the exact reason for Tregs reduction remains unclear, Tregs dysfunction and destabilisation during inflammation may contribute to this observation. However, whether Tregs are unstable during cholangiopathies remains to be investigated as the plasticity of Tregs seems to be organ and context-dependent ^10,11,12^. Besides its main immunoregulatory function, the role of Tregs in modulating stem/progenitor function has been documented in multiple organs such as the skin, brain, and intestines^13–15^. IL2-based approaches to enhance Treg number for treating inflammatory disease have shown promising results in the brain and colon ^16,17^. However, similar approaches are less effective for the liver ^18,19^. Therefore, it is crucial to understand the dynamics of Tregs during liver injury and repair, and whether Tregs have a role in coordinating the bile ducts to undergo compensatory regenerative mechanism during bile duct injury.

Most animal models of experimental cholangitis are limited in recapitulating the complete pathophysiology of PSC to cover both aspects of epithelial tissue damage and the immune landscape of PSC. To determine whether Tregs can influence the regeneration of liver epithelium, we used a combinational approach to investigate the effect of Tregs on DR during experimental cholangitis whilst Tregs are reduced. To further understand the causes of Tregs reduction in PSC, we conducted lineage tracing experiments using a transgenic Foxp3 fate mapper mouse strain to investigate the potential underlying mechanisms of reduced Treg numbers during cholangiopathies. Overall, these experiments highlight potential targets for pharmacological interventions to improve PSC prognosis.

## Results

### Patients with PSC have limited intrahepatic Tregs expansion despite florid ductular reaction

We observed an increase in CD4^+^ T cells in the liver of PSC patients accompanied by an enhanced ductular reaction (DR) (Fig 1A,B). Intrahepatic CD4^+^ T cells are 20-fold higher in livers of patients with PSC compared to livers from healthy non-liver-related death donors (Fig 1B). These intrahepatic CD4^+^ T cells are preferentially localised to the periductal area in PSC livers (Fig 1C). Interestingly, intrahepatic FOXP3^+^ Tregs are not increased in PSC livers compared to healthy livers (Fig 1D). The proximity of CD4^+^ T cells with DR and the limited Tregs expansion in the liver probed us to investigate the relationship between Tregs and bile duct regeneration.

**Fig 1.**
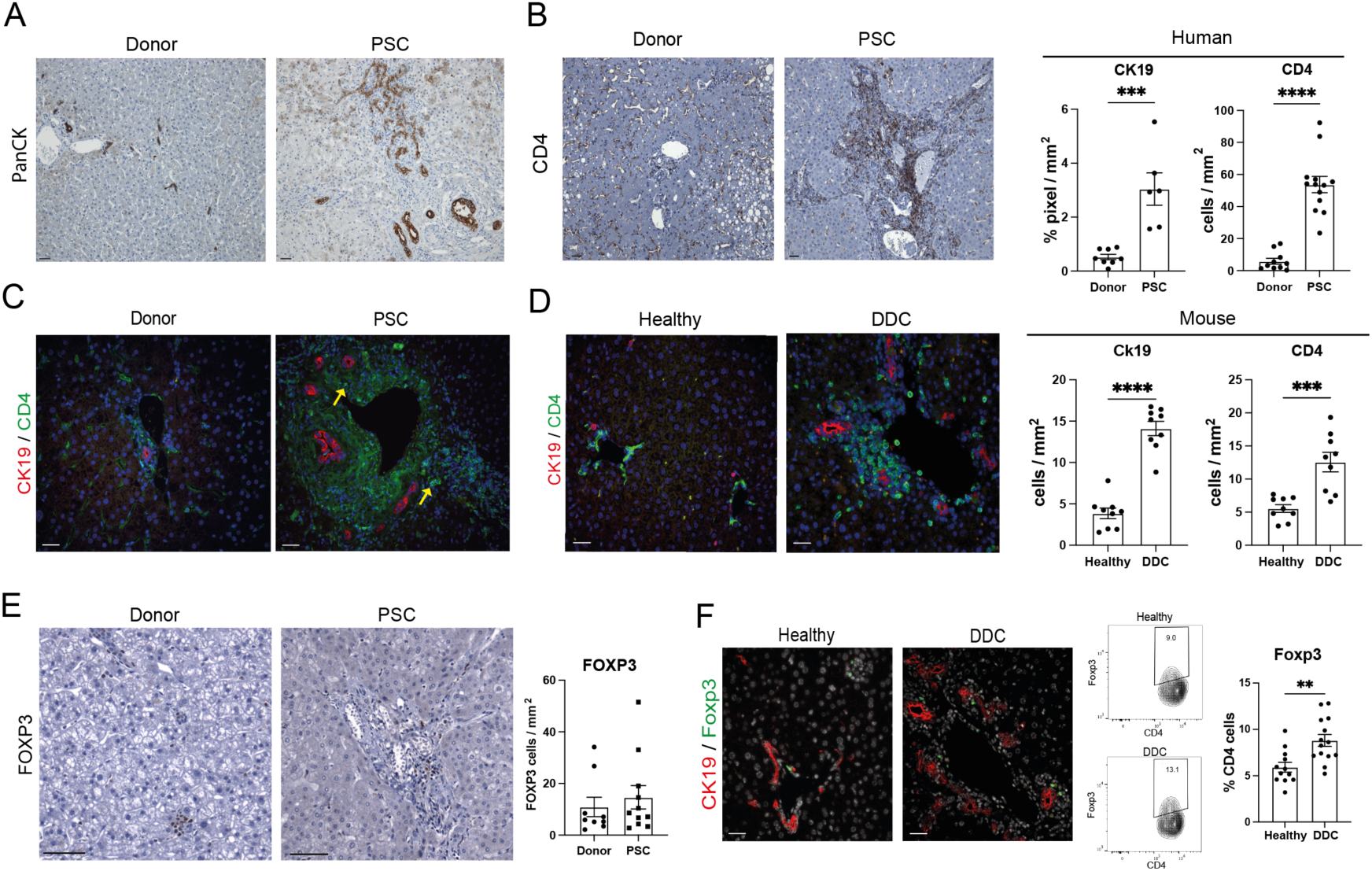
PSC livers have limited FOXP3 Tregs infiltration. Immunohistochemical staining of healthy and PSC liver stained for A) PanCK for cholangiocytes n=8 for healthy, n=6 for PSC B) CD4 for CD4^+^ T lymphocytes and quantification of PanCK for ductular reaction and CD4 for intrahepatic CD4^+^ T lymphocyte infiltration. n=10 for healthy, n=14 for PSC C) Immunofluorescence of CK19 and CD4 on healthy and PSC livers, arrows denote CD4 T lymphocytes near cholangiocytes D) Immunofluorescence of Ck19 and CD4 on healthy and DDC treated mouse livers with quantification of Ck19 for ductular reaction and CD4 for intrahepatic CD4^+^ T lymphocyte infiltration. n=9 mice E) Immunohistochemistry of FOXP3 on healthy and PSC livers with quantification for intrahepatic FOXP3^+^ regulatory T cells. n=9 for healthy, n=11 for PSC F) Immunofluorescence of Ck19 and Foxp3 on healthy and DDC-treated mouse livers with quantification of CD4^+^Foxp3^+^ intrahepatic Tregs by flow cytometry n=12 mice. Data presented as Means ± SEM, Scale bar, 100*μ*m. \**P* < 0.05 and \*\**P* < 0.01, by unpaired two-tailed Mann-Whitney t-test.

To investigate the role of Tregs on bile duct regeneration, we utilised the 3,5-Diethoxycarbonyl-1,4-Dihydrocollidine Diet (DDC) experimental cholangitis model to induce biliary injury. As expected, an increase in intrahepatic CD4^+^ T cells and DR were observed after DDC diet, which is similar to previous observations in PSC livers (Fig 1E). However, DDC diet-induced bile duct injury leads to a 2-fold increase in intrahepatic Foxp3 Tregs predominantly located close to the DR (Fig 1F). The proximity between Tregs and the DR and the fundamental differences in mouse Tregs infiltration compared to human PSC prompted us to investigate whether Tregs determine the efficiency of bile duct regeneration.

### Reduction in Tregs exacerbates bile duct damage and fibrosis

We next investigated whether the reduction of Tregs infiltration in PSC causes impairments in the bile duct repair mechanisms. Tregs have a crucial role in regulating immune responses, with the mutation of Foxp3 or complete ablation of Tregs in mouse models such as in the Foxp3^sf^ scurfy mice causing symptoms of systemic autoimmunity ^20^. To mimic the Tregs phenotype observed in PSC patients, we used the Foxp3^GFPDTR^ transgenic mice which express a knocked-in human diphtheria toxin (DT) receptor and an enhanced GFP(eGFP) gene downstream of the Foxp3 locus which enables us to alter the level of Tregs in a controlled manner through the injection of DT ^21^. The administration of DT transiently reduces Tregs in a dose-dependent manner in these mice (Fig 2A), with Tregs recovering to the normal range within 5 days after DT injection (Supplementary Fig1).

**Fig 2.**
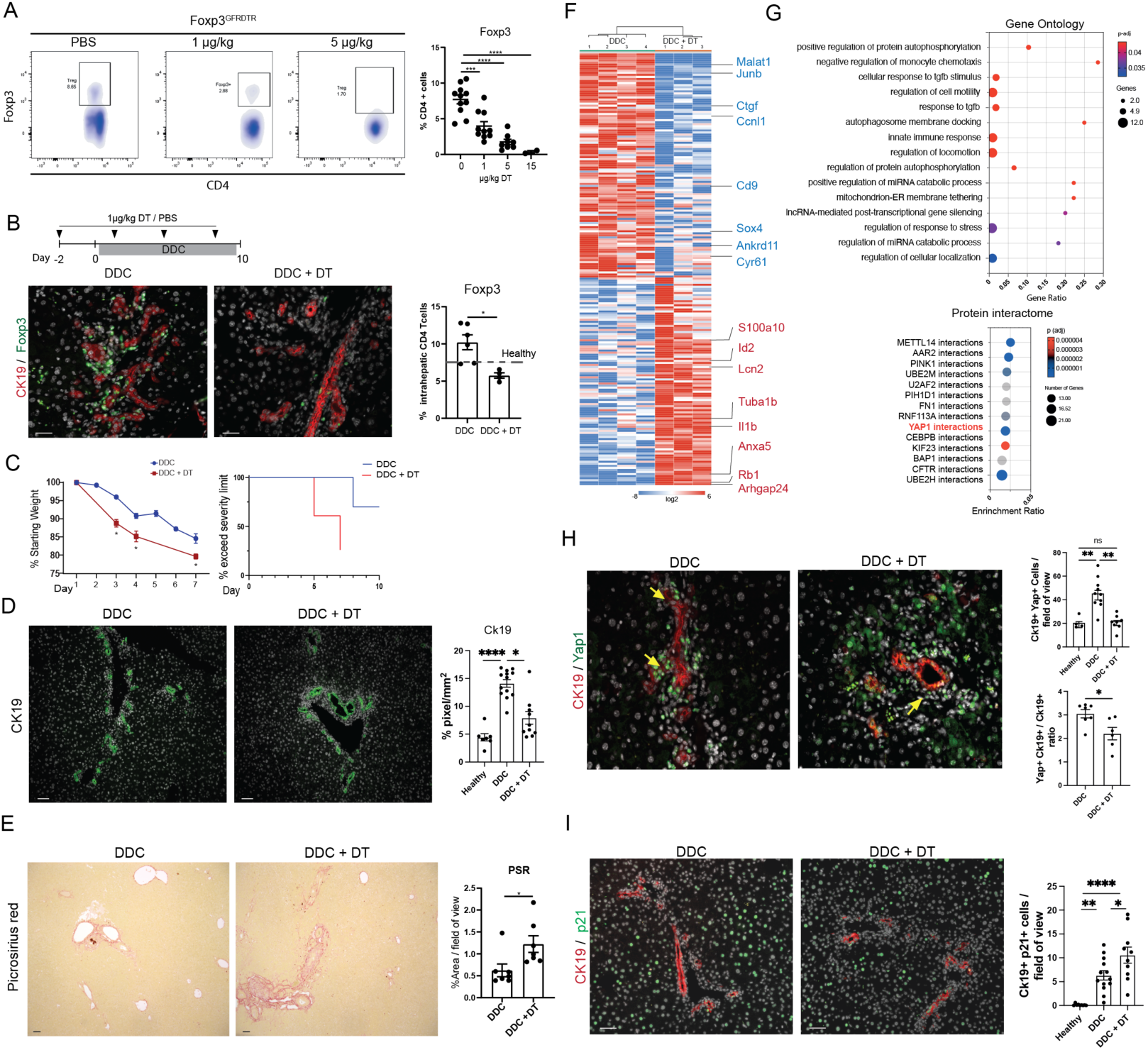
Reduction in Tregs impairs bile duct regeneration and enhances biliary fibrosis. A) Flow cytometry analysis of intrahepatic T cells with quantification of Tregs. *n*=10 mice for 0,1,5 *u* g/kg; *n* = 3 mice for 15 *u* g/kg B) Experimental scheme of DT-mediated intrahepatic Treg depletion on Foxp3^GFPDTR^ mice treated with DDC diet and DT/PBS injection, with dual immunofluorescent staining for CK19 to identify ductular cells and Foxp3 to identify Tregs with quantification of intrahepatic CD4+Foxp3+ Tregs. *n* = 5 mice. C) Graphs showing weight loss and severity level comparing the control and Tregs reduced groups. D) CK19+ cells with quantification showing the magnitude of ductular reaction. *n* = 10 mice. E) Picrosirius Red staining with quantification of Control and DT-treated livers. *n* = 7 mice. F) Heatmap of differentially expressed genes(DEG)genes of EpCAM+ cholangiocytes isolated from Foxp3^GFPDTR^ mice on DDC diet with DT/PBS injection. G) Gene ontology analysis of the top 50 DEG (above), predicted protein interactomes of the top 50 DEG (below) between the control and Tregs reduced mice. H) Representative images of Yap expressing cholangiocytes with quantification of CK19 and Yap1 double positive cells. *n* = 10 mice I) Immunofluorescence staining of CK19 and p21 with quantification of CK19+p21+ senescing cholangiocytes. *n* = 10 mice. Data presented as Means ± SEM, Scale bar, 100*μ*m. \**P* < 0.05 and \*\**P* < 0.01, by unpaired two-tailed Mann-Whitney t test.

By combining the DT injection to the Foxp3^GFPDTR^ mice with the injury-inducing DDC diet regime, we investigate the effect of reduced Tregs during bile duct injury to mimic the lack of Tregs increase in human PSC livers (Fig 2B). Repeated injection of 1μg/kg DT during the DDC diet regime reduced intrahepatic Tregs to the homeostatic level, causing reduced Tregs localised to the periductal area during biliary injury compared to the PBS-injected control group (Fig 2B). Interestingly, mice with reduced Tregs lost weight more rapidly than those with unmanipulated Treg infiltration. Furthermore, the DT-injected group had a lower tolerance to the DDC diet and had to be terminated earlier as they reached the experimental severity limit earlier due to weight loss (Fig 2C). With reduced Tregs infiltration, we observed a 50% reduction in the Ck19^+^ cells in the DT-injected group, suggesting a weaker DR (Fig 2D). The weaker DR was also accompanied by enhanced collagen deposition and fibroblast activation in the DT-injected group (Fig 2E, Supplementary Fig 2D). We performed bulk transcriptomic analysis and compared the transcriptomic changes on EpCAM+ cholangiocytes isolated from mice with intact or reduced intrahepatic Tregs, and identified 210 differentially expressed genes (DEG), (Fig 2F, Supplementary Fig 2A). Gene ontology analysis of the top 50 significant DEG indicates that biological processes including cell motility, Tgfb responses, and immune regulation are affected (Fig2G). Interestingly, interactions with Yap1 were identified as one of the key interactome changes of EpCAM+ cells when intrahepatic Tregs are reduced (Fig2G, Supplementary Fig 2B). This is confirmed with immunohistochemistry showing that the number of Ck19 cholangiocytes that express Yap1 and its downstream target Sox9 are decreased in animals with reduced Tregs (Fig 2H, Supplementary Fig 2C). Furthermore, reduced Tregs also leads to an increased in p21 expressing Ck19 positive senescing cholangiocytes in the liver (Fig 2I). These data highlight the importance of Tregs around the regenerative biliary niche, and reduced Tregs have consequences on the ductular reaction, bile duct integrity and fibrosis.

### Tregs expansion through the IL2 complex has limited effects on improving bile duct regeneration

We then investigate whether increasing the Tregs number by administrating Treg targeting IL2-complexes (IL2c) *in vivo* is a therapeutic option to improve bile duct regeneration. During homeostasis, IL2c administration selectively expand Tregs by three-fold, alongside an increase in Tregs: T effector ratio (Fig 3A, Supplementary Fig 2). A similar trend of Tregs expansion is observed when biliary injury is induced following IL2c administration (Fig 3B,C,D). Surprisingly, despite the increase in intrahepatic Tregs, IL2c administration decreased the magnitude of DR by two-fold (Fig 3E). Furthermore, collagen deposition doubled in the IL2c injected group, with increased serum bilirubin and transaminase levels indicating a worsened liver function (Fig3F, G). Interestingly, we also observed an increase in T-bet+ Th1 cells suggesting that IL2c administration enhances liver inflammation (Fig 3H,I). This is accompanied by an increase in Iba1+ macrophages including Gpnmb+ scar-associated macrophages and a decrease in the anti-inflammatory CD206+ macrophage population (Fig 3J).

**Fig 3.**
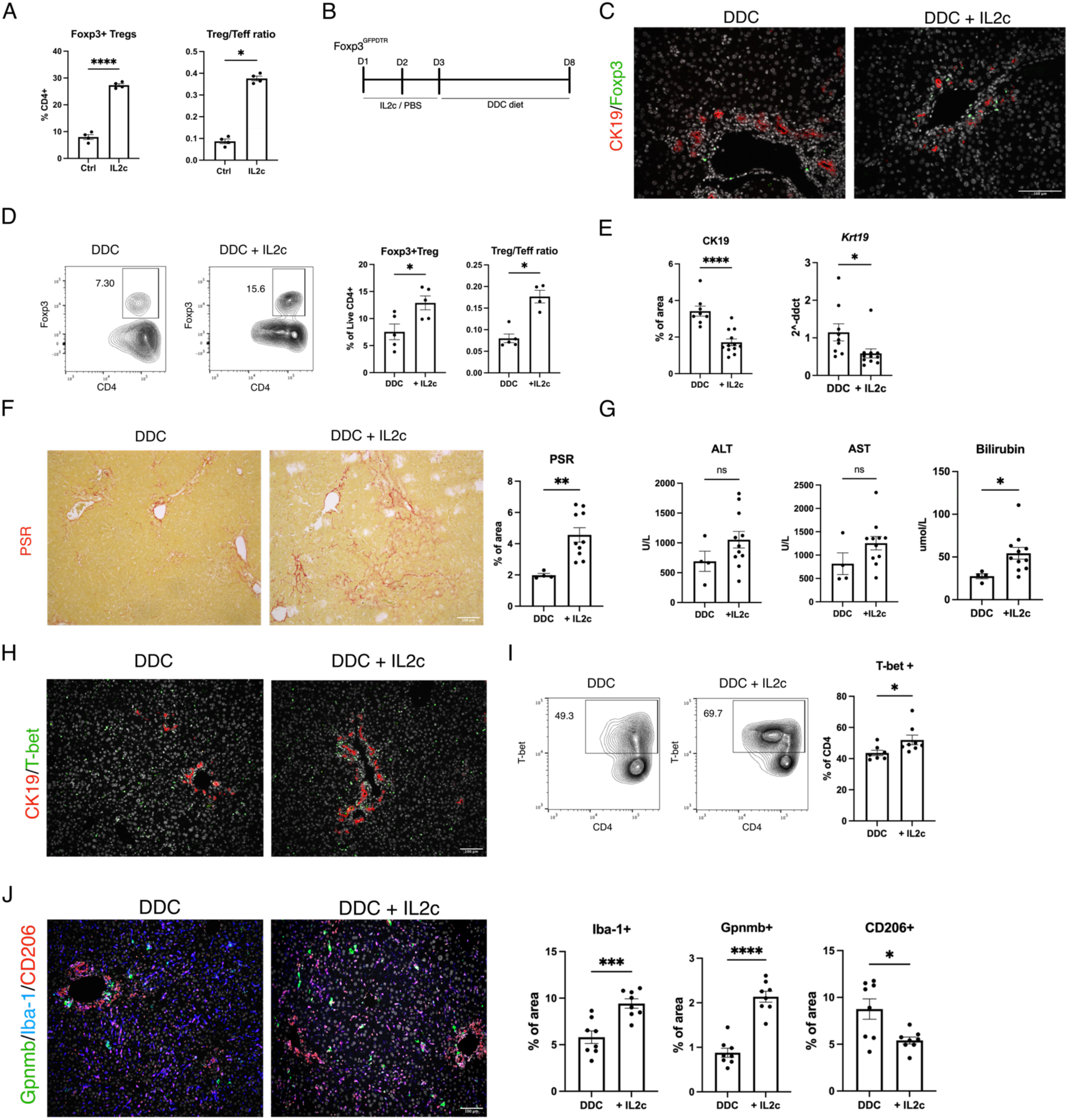
IL2c injection exacerbates bile duct injury. A) Quantification of intrahepatic Foxp3+ Tregs and Tregs (CD4+Foxp3+) to Teff (CD4+Foxp3-) ratio following IL2c injection. B) Experimental scheme of IL2c-mediated intrahepatic Treg expansion during biliary injury C) Double immunofluorescence staining of Ck19 and Foxp3 on DDC-treated livers with and without IL-2c administration. D) Flow cytometry analysis of intrahepatic CD4^+^Foxp3^+^ Tregs and Tregs to Teff ratio. *n* = 5 mice E) Ck19 quantification on DDC-treated livers with and without IL-2c administration and *Krt19* whole liver mRNA expression F) Picrosirius Red staining and quantification of scar area G) Serum liver transaminases and bilirubin levels of treated mice H) Double immunofluorescence staining of CK19 and T-bet. (I) Flow cytometry analysis and quantification of intrahepatic CD4+ T-bet+ Th1 cells J) Immunofluorescence staining of Gpnmb, CD206 and Iba-1 macrophages, with quantifications. Data presented as Means ± SEM, Scale bar, 100*μ*m. *n* = 7-8 mice per group unless specified \**P* < 0.05 and \*\**P* < 0.01, by unpaired two-tailed Mann-Whitney t-test.

### Tregs lose Foxp3 expression during biliary injury and acquire a pro-inflammatory phenotype

We further investigate the underlying cause of the ineffective regeneration in the forementioned IL2c-mediated Tregs expansion experiment and whether this is linked to the reduced Tregs observed in PSC. To understand the dynamic of Tregs during biliary injury, w crossed the Foxp3^GFPCreERT2^ mice with the tdTomato^loxSTOPlox^ mice to generate the Foxp3^GFPCreERT2^tdTomato^loxSTOPlox^ mice, termed the Foxp3^Cre^Ai14 mice hereon. This strain enables us to label Foxp3 Tregs, and investigate the dynamics of Tregs infiltration, proliferation, and stability following biliary injury (Fig 4A). In this strain, Foxp3 Tregs express GFP, and tdTomato expression in Foxp3-expressing cells can be induced upon tamoxifen administration (Fig 4A). With a labelling efficiency of up to 90% of intrahepatic Tregs, we labelled Foxp3 Tregs during homeostasis and investigated the fate of intrahepatic Tregs following DDC-induced biliary injury (Supplementary Fig 4A, Fig 4A). During homeostasis, intrahepatic Tregs are relatively stable with most of the intrahepatic Tregs consisting of labelled Tregs which express Foxp3 and tdTomato. Interestingly, there is a significant increase in the tdTomato-labelled population which does not express Foxp3 following DDC-induced biliary injury (Fig 4B). This indicates that the tdTomato labelled, Foxp3 negative cell population is derived from the Foxp3-positive Tregs, suggesting the loss of Foxp3 expression, we termed this tdTomato^+^Foxp3^-^ population exTreg. Besides the emergence of the exTreg population in the liver, we observed the infiltration of an unlabelled, tdTomato-Foxp3+ Tregs to the liver as expected (Fig 4B). The exTreg population has a lower CD25 expression than Foxp3^+^ Tregs. Upon stimulation, the exTreg population express higher levels of inflammatory cytokines including Ifng and Tnfa (Fig 4C). Furthermore, the exTreg population contributes up to 20% of Tnfa expressing CD4 T-cells during liver injury, suggesting the potential pathological role of the exTreg population (Fig 4D).

**Fig 4.**
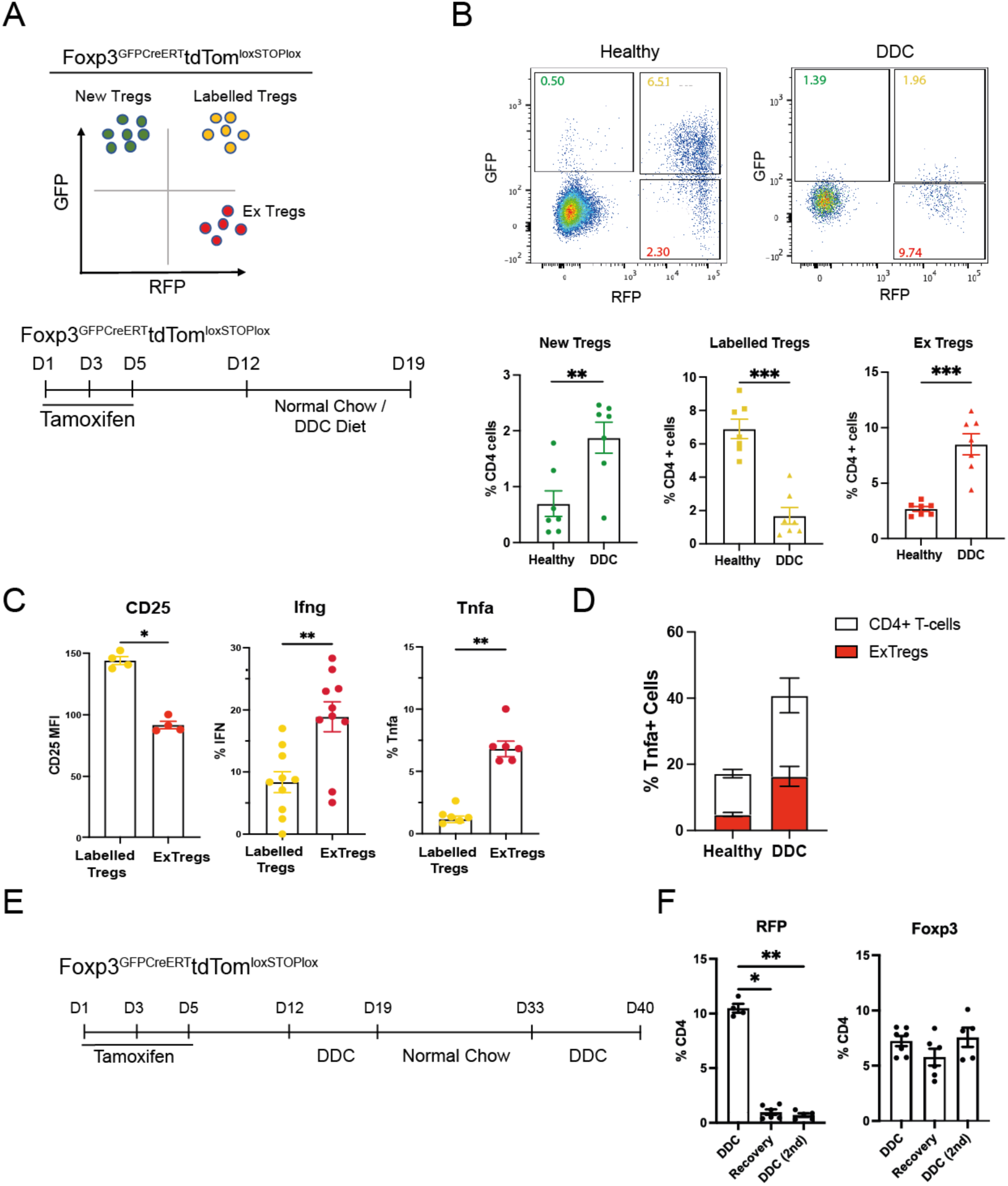
Intrahepatic Foxp3 Tregs acquire a pro-inflammatory phenotype during bile duct injury. A) Flow cytometry gating strategy to identify intrahepatic New Tregs (Green), Labelled Treg (Yellow) and ExTreg (Red) and experimental scheme B) Flow cytometry analysis and quantification of Treg populations. *n*=7 mice C) Flow cytometry analysis of CD25, Ifng and Tnfa expression in labelled and ExTreg. *n*= 4 -10 mice D) Quantification of Tnfa+ cells in healthy and DDC treated mice. *n*=6 mice E) Experimental scheme of long-term Treg fate mapping F) Quantification of intrahepatic CD4+ RFP+ labelled Tregs and Foxp3+ Tregs *n*=6 mice. Data presented as Means ± SEM. \**P* < 0.05 and \*\**P* < 0.01, by unpaired two-tailed Mann-Whitney t-test.

To further investigate whether the exTreg population acquired tissue-resident phenotype, we performed long-term fate mapping experiment post biliary injury. Tregs were labelled in the Foxp3^Cre^Ai14 mice, and the mice were subjected to two rounds of DDC-induced liver injury with a two-week recovery period in between (Fig 4E). Interestingly, we observed that most labelled Tregs are not retained in the liver when injury regressed, and the ones that remain in the liver do not undergo local expansion when the liver is re-exposed to injury. These findings suggest that most intrahepatic Tregs do not originate from the pre-existing Foxp3 population during biliary injury, but derived from newly generated Tregs (Fig 4F). Furthermore, the liver microenvironment may contribute to Treg instability as we only observed the exTreg population in the liver, and limited expression in other organs such as the spleen and the intestines (Supplementary Fig 3B).

### IL2 complexes induce intrahepatic Tregs to acquire a pro-inflammatory phenotype

We then used the Foxp3^Cre^Ai14 mice mentioned above to investigate whether the switch in Tregs stability is the underlying cause of increased fibrosis and inflammation observed in the IL2c-induced Tregs expansion experiments (Fig 5A). Despite the expected Tregs expansion following IL2c administration, fate mapping analysis of Treg populations showed an enhanced emergence of the pro-inflammatory exTreg population compared to the control group which had DDC-induced injury only (Fig 5B,C). We further investigate whether the increase in Tbet+ Th1 population is derived from the exTreg population, and observed that the IL2c administration doubled the tdTomato^+^ Tbet^+^ exTreg population, and no change was observed in the tdTomato-Tbet+ population (Fig 5D,E). This suggests that the IL2c expanded Tregs undergo fate change with the loss of Foxp3 expression and acquire Tbet expression for a pro-inflammatory Th1 phenotype.

**Fig 5.**
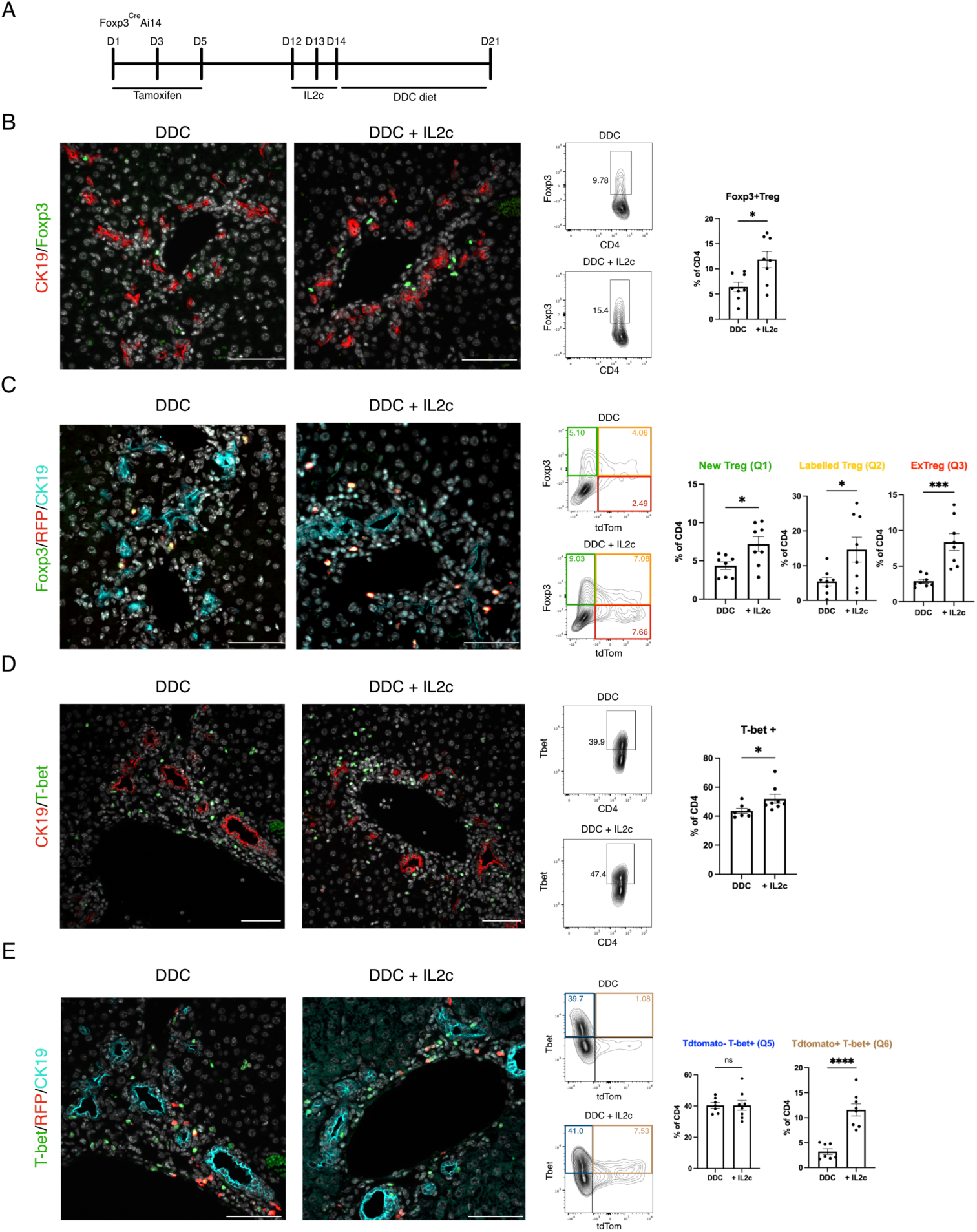
IL2c injection contributes to Tregs plasticity. A) Experimental scheme of IL2c-mediated intrahepatic Treg expansion B) Immunofluorescence staining of CK19 and Foxp3 with flow cytometry analysis of intrahepatic CD4+ T cells with the quantification of intrahepatic Foxp3+ Tregs C) Immunofluorescence staining for Foxp3, RFP and CK19, flow cytometry analysis of intrahepatic CD4+ T cells with the quantification of Treg subpopulations D) CK19 and T-bet, flow cytometry analysis of intrahepatic CD4+ T cells with the quantification of T-bet+ cells E) T-bet, RFP and CK19, flow cytometry analysis of intrahepatic CD4+ T cells with the quantification of T-bet+ subpopulations. Data presented as Means ± SEM. \**P* < 0.05 and \*\**P* < 0.01, by unpaired two-tailed Mann-Whitney t-test.

### Blocking Ox40 signalling alleviates biliary fibrosis

The Ox40 signalling pathway plays an important role in Tregs regulation and stability ^22,23^. We found that intrahepatic Ox40L expression is increased in PSC livers and the expression of Ox40L positively correlates with the degree of scar deposition (Fig 6A). The increase in Ox40L expression is also observed in the mouse DDC injury model, with Ox40L expressed by multiple cell types including macrophages and T lymphocytes (Fig6B, Supplementary Fig4). Furthermore, the Ox40 receptor is predominantly on CD4 lymphocytes, with its expression enriched in the Foxp3+ Treg population (Fig 6C), and the number of Ox40-expressing CD4 T-cells increases during biliary injury (Fig 6D).

**Fig 6.**
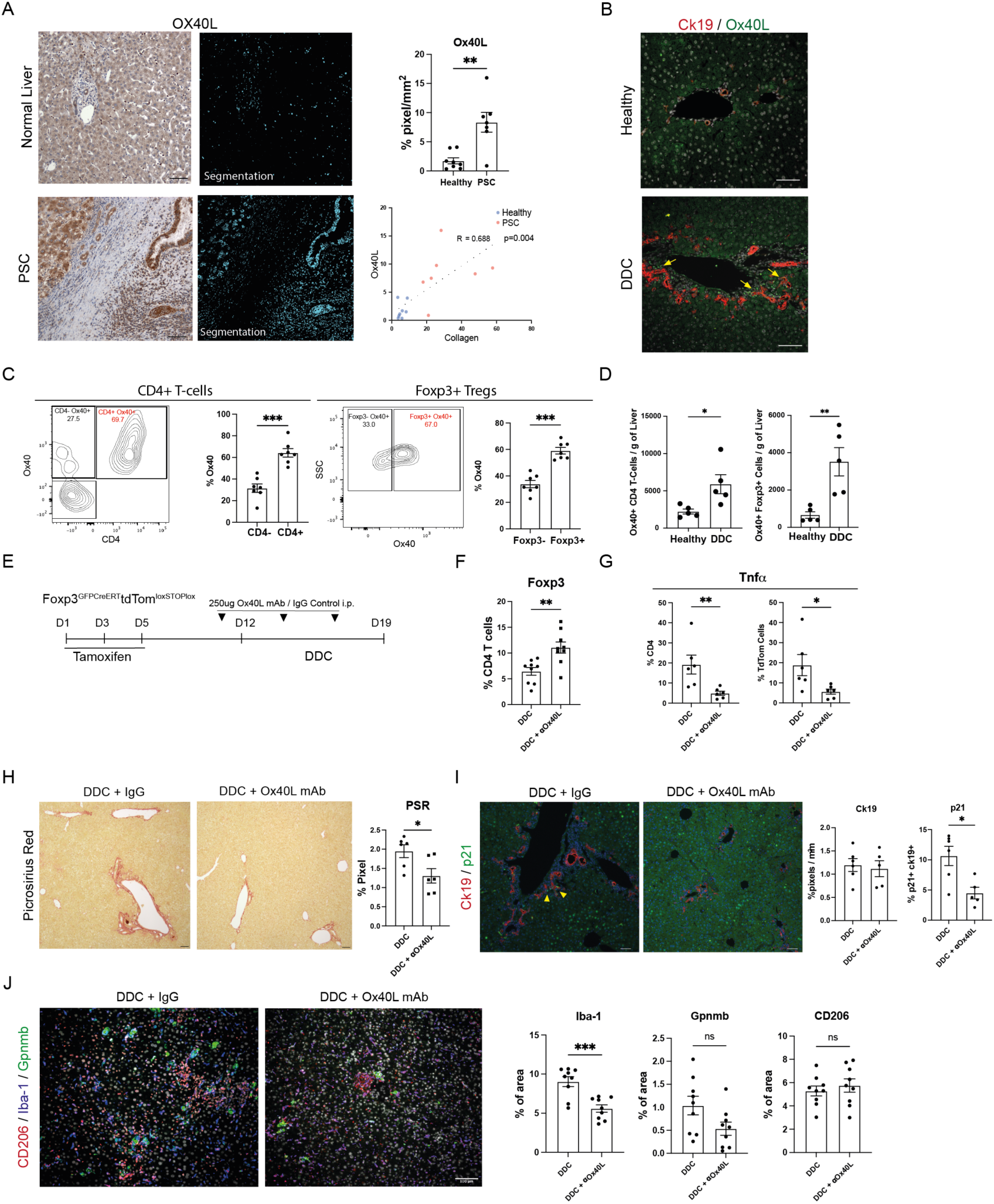
Blocking Ox40L reduces liver inflammation and fibrosis. A) Immunohistochemistry staining of OX40L with quantification in healthy and PSC human liver, correlation curve between liver Ox40L expression and collagen deposition in the liver, *n*=8 for healthy, *n*=7 for PSC B) Representative images of immunofluorescence staining for CK19 and Ox40L (arrows) in healthy and DDC mice C) Flow cytometry analysis of Ox40-expressing cells within intrahepatic CD4+ T-cells and Foxp3+ Tregs, *n*=7 mice. D) Number of intrahepatic Ox40+ expressing CD4+ T-cells and Foxp3+ Tregs per gram of liver in healthy and DDC treated mice. *n*=5 mice E) Schematic showing the experimental design to block Ox40L during the biliary injury. Quantification of intrahepatic Foxp3+ Tregs in control and Ox40L blockade mice post DDC injury. *n*=9 mice G) Tnfa expression post *in vitro* stimulation of isolated intrahepatic CD4+ T-cells and TdTomato+ cells. *n*=6 mice H) Picrosirius Red staining of control and Ox40L blockade mice following DDC induced injury. *n*=6 mice I) Representative images of Ck19 ductular cells and p21 senescence cells with quantification of Ck19+ ductular reaction and CK19+p21+ senescing ductular cells. *n*=6 mice, yellow arrowheads denote senescing cholangiocytes J) Representative images showing macrophage subpopulations in the liver of control and Ox40L blockade mice following DDC induced liver injury. Quantification of Iba-1, Gpnmb and CD206 positive cells in the liver. *n*=9 mice. Data presented as Means ± SEM, Scale bar, 100*μ*m. \**P* < 0.05 and \*\**P* < 0.01, by unpaired two-tailed Mann-Whitney t-test.

We further investigate whether the blockade of Ox40 signalling through the administration of Ox40L blocking antibodies will improve the outcome of bile duct injury and regeneration (Fig 6E). The blockade of Ox40 signalling in the fore-mentioned Foxp3CreAi14 Tregs fate mapping mice increases intrahepatic Tregs number during DDC-induced injury, furthermore, this is accompanied by reduced Tnfa secretion by intrahepatic CD4 T-cells (Fig 6F,G). Moreover, the reduction in Tnfa is also observed in the labelled tdTom exTreg population, suggesting a decrease in liver inflammation and improved Tregs stability (Fig 6G). The blockade of Ox40 signalling also reduced the degree of periportal fibrosis (Fig 6H). However, the increase in Tregs number does not affect the magnitude of Ck19+ ductular reaction and ductular proliferation (Fig 6I, Supplementary Fig 6). Nevertheless, a significant decrease in p21+ senescent cholangiocytes is observed suggesting that the bile ducts are more resistant to cellular senescence when Ox40 signalling is blocked during bile duct injury (Fig 6I). Besides, blocking Ox40L signalling also leads to an increase in total Iba-1 macrophage infiltration with a trend of decrease in the scar-associated Gpnmb^+^ macrophages (Fig6J). Altogether, these data suggest that modulating Ox40 signalling has both immunomodulatory effects and protective effects on bile ducts.

### Blocking Ox40L enhances the efficacy of IL2-mediated Tregs expansion and promotes biliary regeneration

We then explore whether the combination of Ox40L blockade and the administration IL2c promotes bile duct regeneration through increasing Tregs number and stability (Fig 7A). Interestingly, the combination of IL2c and Ox40L blockade enhances the Treg expansion effect of IL2c administration by 2-fold (Fig 7B). The increase in the labelled Treg population and the decrease of the pro-inflammatory exTreg population suggest that more Tregs retain Foxp3 expression when the Ox40 signalling is blocked during IL2 mediated Treg expansion *in vivo* (Fig 7B). Despite the decrease in the exTreg population, no change is observed in the Tbet+tdTomato+ population indicating that blocking Ox40L alone does not fully prevent Tregs from acquiring a Th1 like pro-inflammatory phenotype (Fig 7C). However, the combined effect of Ox40l blockade and IL2c administration increased the Treg: T effector ratio by two-fold compared to IL2c administration alone (Fig 7D). With the change observed in the Tregs, we also observed an increase in Ck19+ DR in the group that received the combination treatment with a decrease in senescing ductular cells and periportal fibrosis (Fig 7E,F,G). The co-administration of IL2c and ox40L blocking antibodies also decreases macrophage infiltration into the liver, and more interestingly, result in a decrease in the Gpnmb^+^ scar-associated macrophage population (Fig 7G). Altogether, these experiments suggest that Tregs can be used as a therapy option to promote bile duct regeneration and decrease periductal fibrosis during cholangiopathies. Furthermore, the efficacy of Tregs-based therapy can be enhanced by targeting co-stimulatory pathways such as the Ox40 signaling.

**Fig 7.**
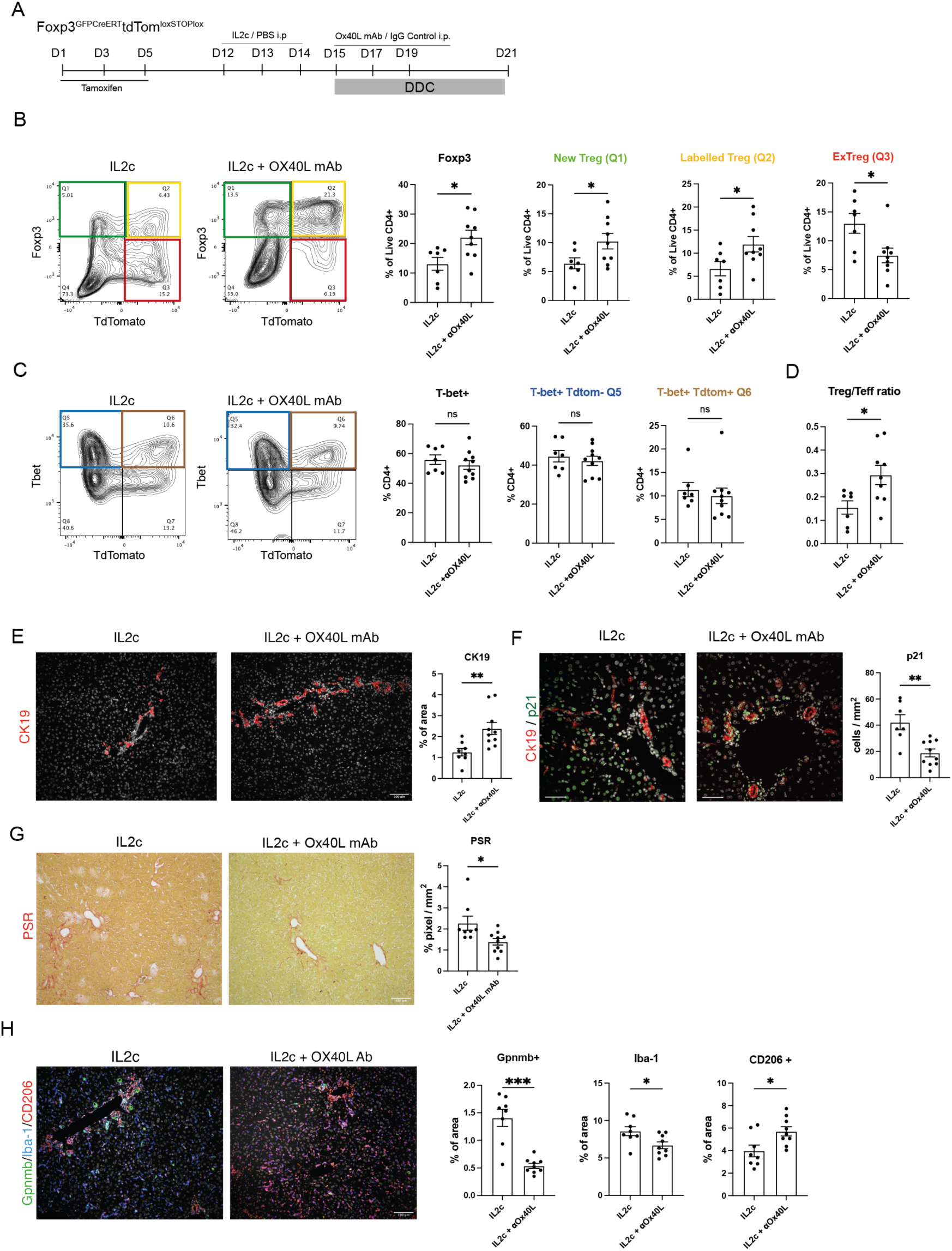
The combination of IL2-complex and Ox40L blocking antibodies enhances Treg numbers and stability. A) Experimental scheme of co-injecting Il2c and Ox40L to expand Tregs during biliary injury B) Flow cytometry gating strategy and quantification of intrahepatic Foxp3+ Tregs and subsequent subpopulations including TdTomato-Foxp3+ New Tregs (Q1,green), TdTomato+Foxp3+ labelled Tregs (Q2, yellow), TdTomato+Foxp3-ExTreg (Q3, red) in IL2c injected and IL2c + Ox40L antibodies co-injected mice. (*n* =7-10 mice) C) Flow cytometry analysis of intrahepatic CD4+ T cells with the quantification of CD4+T-bet+ populations and subsequent subpopulations including conventional Th1 cells, Tbet+Tdtomato-cells (Q5), Tregs derived Th1 like cells, Tbet+TdTomato+cells (Q6) *n* =7-9 mice D) Intrahepatic Tregs to T effector ratio. E) Immunofluorescence staining of Ck19 ductular cells in mice treated with IL2c or IL2c + ox40L antibodies. *n*=10 mice F) Representative images showing Ck19 ductular cells and p21 as a marker of cellular senescence with quantification of CK19+p21+ double positive cells. *n*=10 mice G) PSR staining for collagen deposition in the liver with quantification. *n*=10 mice H) Representative images showing macrophage subpopulations in the liver of IL2c injected and IL2c + Ox40L blocking antibodies co-injected mice post DDC induced liver injury. Quantification of Iba-1, Gpnmb and CD206 positive cells in the liver. *n*=8-9 mice. Data presented as Means ± SEM, Scale bar, 100*μ*m. \**P* < 0.05 and \*\**P* < 0.01, by unpaired two-tailed Mann-Whitney t-test.

## Discussion

The underlying cause of PSC remains unknown, however, recent evidence pinpoints the pathobiology of PSC as immune-triggered ^6,24^. The disease mechanisms of PSC have been compared to other chronic liver diseases, especially PBC in most instances due to their similarities in causing cholangiopathies and both being T-cells driven. However, fundamental differences such as sex prevalence, the location of bile duct damage, and differences in immune cell composition between PSC and PBC may explain the difference in the effectiveness of treatments such as ursodeoxycholic acid (UDCA) therapy ^25^. Interestingly, reduced number of Foxp3 Tregs has been observed in PSC compared to PBC ^9,18^. Tregs promote tissue repair in multiple organs by interacting with tissue stem/progenitor cells besides modulating the immune responses surrounding the regenerative niche. Whether Tregs promote epithelial regeneration in the liver remains to be investigated, especially in the context of immune-mediated bile duct damage ^13,15^. Existing animal models of experimental cholangiopathies are limited in recapitulating the distinct Tregs phenotype in livers of patients with PSC^26^, and this hinders the understanding of disease mechanism and development of efficient therapies albeit great advances in Genome-wide association studies (GWAS) about PSC in the past decade^27,28^. Tregs-based therapies remain a popular therapeutic option for PSC since their beneficial effects are shown in organs such as the skin, muscles and bones^29,30^. On the contrary, the efficacy of Treg-based therapies are less promising in the liver with further optimisation required^18,19,31^.

To understand the underlying mechanisms and improve the efficacy of Treg-based therapy for cholangiopathies, we combined animal models of immunomodulation and experimental cholestasis to understand whether reduced intrahepatic Tregs affect the bile duct regenerative responses to injury. We mimicked the lack of intrahepatic Tregs expansion in PSC livers using the dose-dependent diphtheria toxin inducible Foxp3^GFPDTR^ Tregs depletion model to reduce Treg number in the liver without triggering systemic autoimmunity ^21^ ^32^. Interestingly, the reduction of Tregs around the periductular niche during bile duct injury reduced the magnitude of ductular reaction (DR) but increased periductular fibrosis. The inverse correlation between DR and fibrosis we observed contradicts the conventional thought which suggests that DR leads to the development of fibrosis ^33,34^. Transcriptomic analysis performed on isolated bile ducts suggests that Tregs regulate the DR and bile duct integrity in response to injury, with pathways involved in DR such as Yap and Tgfb responses affected when Treg numbers are reduced. The dysregulation in these pathways leads to decreased bile duct integrity and promotes biliary senescence ^35,36^. Hence, the lack of Tregs infiltration observed in PSC livers may be one of the underlying causes of increased cholangiocyte senescence observed in PSC ^37,38^. In the series of experiments, which we reduced and expanded Tregs, the magnitude of DR does not always correlate directly with the degree of fibrosis, indicating that the link between DR and fibrosis can be uncoupled. Tregs are the key regulators that determine how cholangiocytes respond to injury whilst exerting its immunomodulatory effect on the periductular niche, highlighting the importance of Tregs in regulating the consequences of DR, uncoupling the long existing dogma regarding the pathogenic and pro-fibrotic effect of DR.

We also used a transgenic Treg fate-mapping model to investigate the dynamic of Tregs in response to bile duct injury, as factors such as Tregs infiltration, expansion, or stability might be the contributing factors to the lack of Tregs numbers observed in PSC livers^9^. Using a mouse model of experimental cholangitis, we observed that Foxp3^+^ Tregs lose Foxp3 expression and acquire a pro-inflammatory phenotype with higher expression of inflammatory cytokines such as Tnfa and Ifng during injury, which we termed exTreg for this study. Furthermore, most labelled Tregs are not retained in the liver, and those that are retained do not undergo local expansion, suggesting the transient tissue residency phenotype as shown recently^39^. The acquisition of the exTregs phenotype during bile duct injury is worth noting as the adoptive transfer of Tregs has been proposed as a potential therapy for liver diseases^40^. The stability of transplanted Tregs and the retention of anti-inflammatory, immunomodulatory function is critical for the success of Tregs-based therapy and remains to be investigated. This is further confirmed by our IL2c Tregs expansion experiments that, despite successfully expanding Tregs using IL2c, the loss of Foxp3 expression and the acquisition of pro-inflammatory phenotype can be detrimental to bile duct regeneration.

We then investigated whether we could address the issue of Tregs instability by targeting immunoregulatory signals. The Ox40 co-stimulatory signal is crucial in regulating immune function and Tregs stability ^22,23^. Upregulation of Ox40L has been observed in several inflammatory diseases, including metabolic dysfunction-associated steatotic liver disease (MASLD), and the blockade of Ox40L signalling reduces fibrosis through remodelling the inflammatory niche ^41,42^. Here, we report the upregulation of Ox40L coincides with fibrosis levels in PSC livers. More importantly, fibrosis level is reduced through Ox40L blocking antibodies administration in experimental cholangitis model. Mechanistically, blocking Ox40L reduced Tnfa secretion suppressed the inflammatory phenotype of the exTreg population and reduced liver macrophage infiltration. However, the blockade of Ox40L signalling alone did not enhance ductular proliferation despite reducing cholangiocyte senescence levels. Besides affecting T-cells and cholangiocytes, the blockade of Ox40L also reduces the infiltration of lipid/scar-associated Gpnmb-expressing macrophages, suggesting that Ox40L blockade affects both the adaptive and myeloid compartments^43^. Blocking Ox40L does not alter the number of CD206 expressing anti-inflammatory macrophages, whether the function of these cells is affected following Ox40L blockade remains to be investigated as CD206 macrophages have been reported to be dysfunctional in PSC patients ^24^. More importantly, combining Ox40L blockade with IL2c administration further enhanced the number of Tregs, reduced the emergence of the pro-inflammatory exTreg population, decreased liver fibrosis and improved bile duct regeneration compared to administrating IL2c alone. However, the incorporation of Ox40L blockade with IL2-c does not completely ameliorate the symptoms of cholangitis. For example, blocking Ox40L prevents Tregs from downregulating Foxp3 but does not prevent the upregulation of Tbet suggesting that the maintenance of Foxp3 expression and the acquisition of Tbet are governed by different mechanisms such as the IL6 and IL12 signalling ^18,44^.

Nevertheless, we have demonstrated the importance of maintaining Tregs stability when utilising Tregs therapy for cholangitis as Tregs stability may be the limiting factor for Tregs-based therapies for liver disease. We also proposed a combinational approach of enhancing Tregs number whilst targeting co-stimulatory signals such as the Ox40 pathway to alleviate symptoms of cholangitis. Undoubtedly, Tregs are an attractive and promising source for treating inflammatory diseases, however, a better understanding of Tregs interaction with the tissue microenvironment is required and should be further explored to increase the efficacy of Tregs-based therapies.

### Limitations of the study

We acknowledge that there is currently no perfect animal model for PSC and our combinational approach to alter Tregs level whilst inducing cholangiocyte damage is still insufficient to fully recapitulate the complex pathology of PSC. Nevertheless, our mouse studies highlight the importance and the potential of Tregs in regulating cholangiocyte-initiated ductular response besides its immunomodulatory effects. The observation that Tregs become phenotypically unstable in a highly inflammatory environment might explain the lack of Tregs infiltration observed in PSC patients as Tregs enter the inflamed liver and lose Foxp3 and other Tregs markers such as CD25. Whether this observation from our mouse studies occurs in humans remains to be investigated. We are aware that our Tregs stability study does inform us whether the unstable Tregs subpopulations originate from natural Tregs or induced Tregs since both natural and induced Tregs express the signature transcription factor Foxp3. However, the origin of Tregs populations in also unclear in cholangitis. The lack of a definitive exTreg marker limits the study of Tregs stability in humans, and further work is required to understand the molecular mechanisms of the transitioning Tregs population and stability.

## Acknowledgements

This work was supported by the UKRI MRC Career Development Award (MR/T030798) (to W.Y.L) and the PSC Partners Seeking a Cure Young Investigator Award (to W.Y.L). We thank the Niigata Foundation Promotion of Medicine support (to N.K), MRC Support of (MR/W006804/1) (to B.H). GA is funded by MRC Programme Grant (MR/T029765/1). The Authors acknowledge the University of Edinburgh IRR Flow Cytometry and Cell Sorting Facility, Biomolecular Core facility, Bioresearch & Veterinary Services of the University of Edinburgh and the University of Birmingham. We thank David Bending, Kendle Maslowski, Sarah Dimeloe, Rebecca Drummond for their advice. We thank the Lothian NRS Bioresource, the Centre for Liver and Gastrointestinal Research Birmingham, and patients who consented and donated tissue for research.

## Funding

UKRI MRC Career Development Award (MR/T030798) (to WYL)

PSC Partners Seeking a Cure Young Investigator Award (to WYL)

Niigata Foundation Promotion of Medicine support (to NK),

MRC Precision Medicine Studentship Support of (MR/W006804/1) (to BH)

MRC Programme Grant (MR/T029765/1) (G.A)

## Author contributions

Conceptualisation: WYL

Investigation: NK, MCW, GH, BH, PCH, MB, DP, TK

Data Analysis: MCW, NK

Funding acquisition: AT, ST, WYL

Supervision: AGR, PR, GA, DW, SS, WYL

Writing – original draft: MCW, NK, WYL

Writing – review & editing: AGR, PR, GA, DW, SS, AT, WYL

## Material and Methods

### Animal studies

All animals were housed and bred in temperature- and light-controlled facilities and were maintained following the guide and use of Laboratory Animal and the Animal Welfare Act. All experiments on animals were approved by the University of Edinburgh and the University of Birmingham’s local ethical committee, and local veterinary guidance and studies were performed under the approval of the UK Home Office project licence PP5982214. Both male and female mice aged from 8-16 weeks were used for these experiments.

Foxp3^GFPDTR^ mutant mice (Jax no: 016958) express knocked-in human diphtheria toxin (DT) receptor and GFP genes from the Foxp3 locus. Mice were injected with Diphtheria Toxin (DT) (Sigma-Aldrich) dissolved in PBS at the concentration of 1, 5 or 15μg/kg depending on the experimental design with PBS used as vehicle control.

Foxp3^CreERT2^Ai14 was generated by crossing the Foxp3^GFPCreERT2^ (Jax no:016961) and Ai14 mice (Jax no: 007914) mice. Tamoxifen(4mg) was dissolved in olive oil and administered through oral gavage 3 times per week.

Mice were harvested according to UK Home Office regulations, blood was collected by inferior vena cava (IVC) puncture and centrifuged at 10000g to collect serum. Analysis of serum was performed by the University of Edinburgh Biomolecular Core facility. Organs were harvested and either directly frozen at −80 °C or fixed in 10% formalin for 12 hours before paraffin embedding.

### 5-diethoxycarbonyl1,4-dihydrocollidine (DDC) diet

To induce biliary injury, mice were given 0.1% 3,5-diethoxycarbonyl1,4-dihydrocollidine (DDC) diet for 6 -10 days. Weight loss of more than 20% of starting weight was used as the experimental humane endpoint.

### Interleukin-2 complex (IL-2c) administration

IL-2c comprised 1μg of murine recombinant interleukin-2 protein (Biolegend) and 5μg of purified anti-mouse IL2 antibody (clone JES6-1A12). IL-2c was diluted in 100 μL of sterile PBS before the intraperitoneal injection. Every mouse received 3 doses of either IL-2c or PBS control consecutive days before the DDC diet. All mice were given the DDC diet and culled on day 5 after the start of the DDC diet.

### OX40L blocking antibodies administration

250 μg of OX40L Ab (InVivoMab, Clone RM134L) in 100 uL of PBS were injected into the mice every other day within the DDC diet period. A total of three doses of OX40L Ab or rat IgG2b control were injected.

### Liver cell isolation and analysis

To isolate intrahepatic lymphocytes, liver tissues were minced and digested in DMEM (Gibco) + 0.01 mg/ml DNase I and 0.1mg/ml Collagenase IV in a shaking incubator at 37°C for 30 min. The suspension was then homogenised in a gentleMacs dissociator (Miltenyi) and passed through a 70µM cell strainer (Greiner bio-one). The cell suspension was centrifuged at 700g for 20 min with 33% Percoll and the cell pellet enriched with intrahepatic lymphocytes was used for further analysis. For the isolation of other liver cell types, mouse liver tissues were minced and digested using Liver Dissociation Kit (130-105-807, Miltenyi) following the manufacturer’s protocol. The final cell suspension was washed in PBS, passed through a 70 µM cell strainer (Greiner bio-one) and centrifuged at 400g for 5 min and the cell pellet was used or further analysis.

### Splenocyte and T-cell isolation

The spleen was gently pushed through a 70uM cell strainer and centrifuged at 400G for 5 min and cell pellets containing solenocytes were used for further analysis. For T-cell isolation, T-cell isolation kits (Biolegend, 480137, 480005) were used to isolate CD4 T-cells and regulatory T-cells for further downstream workflow following manufacturer procedures.

### Flow Cytometry and Sorting

Single cell suspensions were incubated with Zombie NIR cell viability kit (Biolegend, 423105) and washed with PBS. Cells were then incubated with respective FACS antibodies cocktail at appropriate concentrations, reconstituted in PBS with 2% FCS. Antibody panel: Alexa Fluor 700 anti-CD3 (17A2, Biolegend), APC anti-CD4 (GK1.5, Biolegend), BV786 anti-CD45 (30-F11, Biolegend), BV711 anti-F4/80 (BM8, Bio Legend), BV650 anti-NK1.1 (PK136, Bio Legend), BV510 anti-CD11b (M1/70, Bio Legend), PerCP anti-CD8 (53-6.7, Bio Legend), BV421 anti-IL17a (TC11-18H, Bio Legend), FITC anti-Foxp3 (FJK-16s, Invitrogen), PE-cy7 T-bet (Bio Legend), PE anti-CD4 (RM4-5, Bio Legend), and PE-Cy7 anti-OX40L (RM134L, Bio Legend). Analysis was performed on a BD LSR Fortessa II analyser and subsequent analysis was performed with FlowJo Software (BD).

### RNA-Sequencing and Data Analysis

RNA-sequencing (RNA-seq) was performed on FACS sorted CD45-CD31-Ter119-EpCAM+ cholangiocytes from Foxp3^GFPDTR^ mice which received DDC diet with PBS/DT administration (DDC vs DDC+DT; n = 4 and 3 respectively) that were sorted using a FACS Aria II cell sorter (BD Biosciences). Each sorted cells sample had a purity of >90%. RNA extraction was performed using the PicoPure RNA Isolation Kit (Thermo Fisher Scientific) following the manufacturer’s instructions.

Total RNA quality was assessed using the Agilent 2100 Bioanalyzer System. Extracted RNA samples were processed using the QuantSeq 3′ mRNA-Seq Library Prep Kit (Lexogen, Vienna, Austria) and sequenced on an Illumina NextSeq 500. Read counts were normalized for effective library size. Data were analysed using ROSALIND^®^ (https://rosalind.bio/) platform (ROSALIND, Inc., San Diego, CA, USA). QC metrics, such as imaging quality, binding density, positive control linearity and limit of detection, were inspected. The identification of DEGs between specified groups was performed using ROSALIND^®^. The Benjamini-Hochberg method was applied for adjustment of *p*-values to correct for multiple comparisons. Genes with fold change> 1.5 and adjusted *p*-value < 0.05 were defined as DEGs in ROSALIND^®^. Based on *P* values, we determined the top 50 genes that were significantly up- or down-regulated between (DDC vs DDC+DT group). Gene list enrichment was analysed using ToppGene Suite (http://toppgene.cchmc.org)^45^. Heatmaps were generated with MORPHEUS (https://software.broadinstitute.org/morpheus) from data extracted from the ROSALIND platform, protein association networks were analysed with STRING (string-db.org)^46^.

### Human samples

Anonymised unstained FFPE sections of human liver sections were obtained from the Lothian NRS Bioresource (Approval No.REC ref – 20/ES/0061) after approved application. Livers sections of Non-liver-related death and PSC patients were obtained and analysed.

### Immunostaining and Immunofluorescence staining

For immunohistochemistry, 10% formalin-fixed tissue was cut into 4-µm-thick sections. When necessary, heat-mediated antigen retrieval was performed in 10mM sodium citrate buffer at pH 6.0. For 3,3’-diaminobenzidine (DAB) staining, sections were blocked using 3% hydrogen peroxide for 10 minutes at room temperature. Primary antibodies were incubated overnight at 4℃. Species-specific biotinylated antibodies were used for secondary detection. Slides were then stained using a Vectastain ABC kit (Vector Laboratories, Inc., Burlingame, CA). Nuclei were stained using haematoxylin (Vector Laboratories, Inc.).

Formalin-fixed tissue embedded in paraffin 4μm sections were used for immunofluorescence staining. These slides were blocked with protein block (Spring Bioscience) and stained overnight at 4℃ using antibodies listed in the table. Primary antibodies were detected using fluorescent conjugated secondary antibodies (AlexaFluor 488/ 555 / 647; Invitrogen) for 1 hr at RT. Sections were stained with Dapi and mounted with fluromount (SouthernBiotech).

Images were obtained with an Olympus Bx53 Microscope with MicroPublisher 6 camera, 10 – 20 random non-repeats images covering 1mM^2^ were taken per sample.

Quantification was performed using ImageJ software.

### Polymerase Chain Reaction

Liver tissue was homogenised in Trizol (Life Technologies). Homogeneates were mixed with chloroform (1:5 ratio Chloroform: Trizol) and centrifuged 10000g, for 15 minutes. The aqueous supernatant was removed and mixed 1: 1 with 70% ethanol. RNA was extracted using Qiagen RNaesy mini kit according to the manufacturer’s instructions. Total RNA was reverse-transcribed using a Qiagen Quantitect reverse transcription kit (Qiagen) on a LightCycler 480 II (Roche). Gene expression analysis was achieved by using pre-validated QuantiTect primers (Supporting information Table) with QuantiTect SYBR reagent (Qiagen). Gene expression was normalized to the housekeeping gene, *Ppia*. The fold change in relative gene expression from the control was calculated by using the ΔΔCt method.

### Statistics

For comparison between two groups, a two-tailed *t*-test was performed. For analysis of gene expression data between more than two groups, a one-way analysis of variance (ANOVA) was performed. For analysis of cell counts, such as proliferating hepatocytes, a Mann-Whitney U test was performed. For analysis of change in body weight over time, a two-way ANOVA was performed. *P* < 0.05 was considered significant, bar charts are presented as means ± SEM. All statistical analysis and graph generation were performed using GraphPad Prism 7 software.

