## Supplementary Figures for "Regulatory T cell stability determines the efficiency of bile duct regeneration during cholangitis"

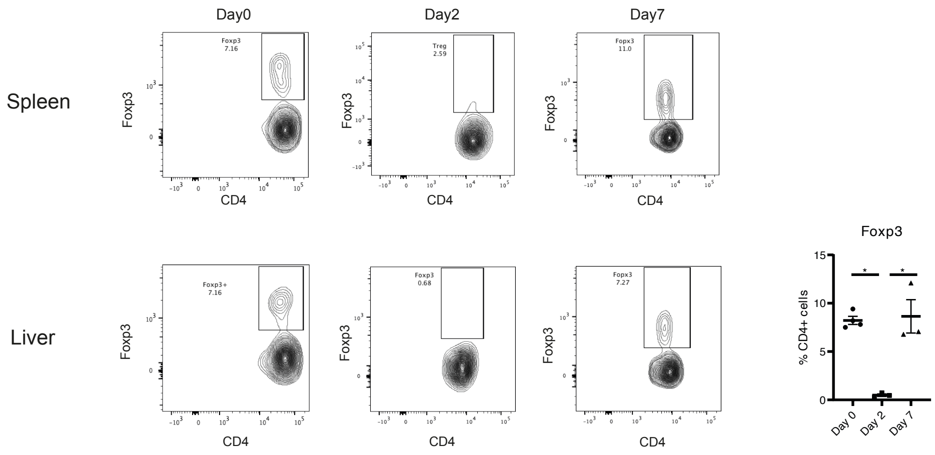


**Supplementary Figure 1. Diphtheria Toxin injection transiently reduces Treg numbers in the Foxp3$^{GFPDTR}$ mice.**

Flow cytometry analysis and quantification of Foxp3+ Tregs in liver and spleen following 5ug/kg injection of Diphtheria Toxin. *n* = 4 mice

A

Top 50 significant change

| Name | Fold Change |
|---|---|
| Erdr1 | -27.9902 |
| Malat1 | -2.05617 |
| Rsrp1 | -3.03696 |
| Dusp1 | -2.05144 |
| Junb | -2.15269 |
| Ctgf | -2.008 |
| Ankrd1 | -1.55173 |
| Anxa5 | 1.77944 |
| Kcnq1ot1 | -2.09046 |
| Kmt2a | -3.59188 |
| Rps20 | 1.96842 |
| Map4k4 | -2.14793 |
| Dbi | 1.7746 |
| Spag9 | -3.31385 |
| Flnb | -1.99009 |
| Ccnl1 | -2.45573 |
| Rpsa | -2.51713 |
| Iqgap1 | -1.86962 |
| 9130008F23Rik | -3.14136 |
| Scara3 | 2.25683 |
| Amotl1 | -2.47926 |
| Mdn1 | -3.03282 |
| Tstd1 | 2.41809 |
| Rap2b | 2.36172 |
| Neat1 | -1.99996 |
| Csrnp1 | -3.52978 |
| Psma3 | 3.19816 |
| Zyx | -2.87954 |
| Cnih1 | -3.67252 |
| Tgm2 | 1.85283 |
| Tceb2 | 3.11107 |
| Nfkbia | -1.56536 |
| Pum2 | -2.60591 |
| Prdx1 | 1.82867 |
| Rps12 | 1.50578 |
| Rab11a | -2.83162 |
| Alb | 1.65558 |
| Lcn2 | 1.89452 |
| Skil | -2.63686 |
| Vmp1 | -1.91751 |
| Zrsr2 | -2.73509 |
| Calm1 | 1.56295 |
| Sdha | 2.90414 |
| Ifrd1 | -1.7573 |
| Hspa1a | -1.94163 |
| Usp7 | -2.27229 |
| Hmgcs1 | 2.37919 |
| Mapk7 | -3.46525 |
| 1190002N15Rik | -2.10678 |
| Srsf11 | -1.52585 |
| Cstb | 1.71901 |

Top 20 significant downregulated

| Name | Fold Change |
|---|---|
| Erdr1 | -27.9902 |
| Malat1 | -2.05617 |
| Rsrp1 | -3.03696 |
| Dusp1 | -2.05144 |
| Junb | -2.15269 |
| Ctgf | -2.008 |
| Ankrd1 | -1.55173 |
| Kcnq1ot1 | -2.09046 |
| Kmt2a | -3.59188 |
| Map4k4 | -2.14793 |
| Spag9 | -3.31385 |
| Flnb | -1.99009 |
| Ccnl1 | -2.45573 |
| Rpsa | -2.51713 |
| Iqgap1 | -1.86962 |
| 9130008F23Rik | -3.14136 |
| Amotl1 | -2.47926 |
| Mdn1 | -3.03282 |
| Neat1 | -1.99996 |
| Csrnp1 | -3.52978 |

Top 20 significant upregulated

| Name | Fold Change |
|---|---|
| Anxa5 | 1.77944 |
| Rps20 | 1.96842 |
| Dbi | 1.7746 |
| Scara3 | 2.25683 |
| Tstd1 | 2.41809 |
| Rap2b | 2.36172 |
| Psma3 | 3.19816 |
| Tgm2 | 1.85283 |
| Tceb2 | 3.11107 |
| Prdx1 | 1.82867 |
| Rps12 | 1.50578 |
| Alb | 1.65558 |
| Lcn2 | 1.89452 |
| Calm1 | 1.56295 |
| Sdha | 2.90414 |
| Hmgcs1 | 2.37919 |
| Cstb | 1.71901 |
| Rpl14 | 2.10564 |
| Cox4i1 | 1.95325 |
| Cox7a2 | 2.20571 |

B

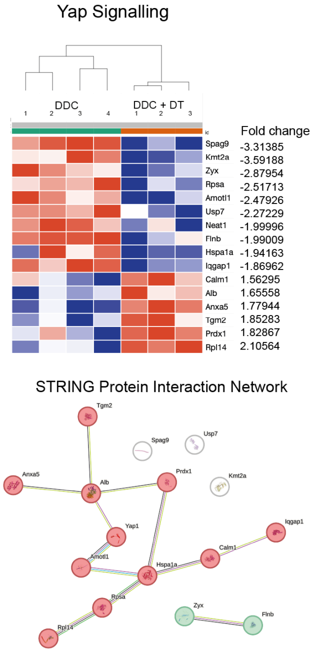


C

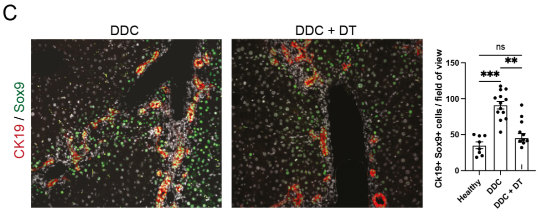


D

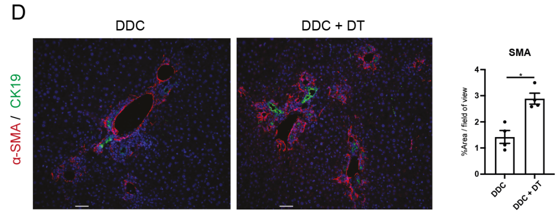


**Supplementary Figure 2. List of top differentially expressed genes following Tregs reduction.** A) Gene list of top 50 significant change (left), top 20 significant downregulated (middle), top 20 significant upregulated (right). B) Heatmap of genes amongst the top 50 DEG which are predicted to be interacting with Yap1 (top), with STRING protein interaction network visualisation (bottom). C) Representative images with quantification of Ck19+Sox9+ double positive cells. *n* = 10 mice D) Double immunofluorescence staining of CK19 and a-SMA with quantification of a-SMA+ area *n* = 4 mice.

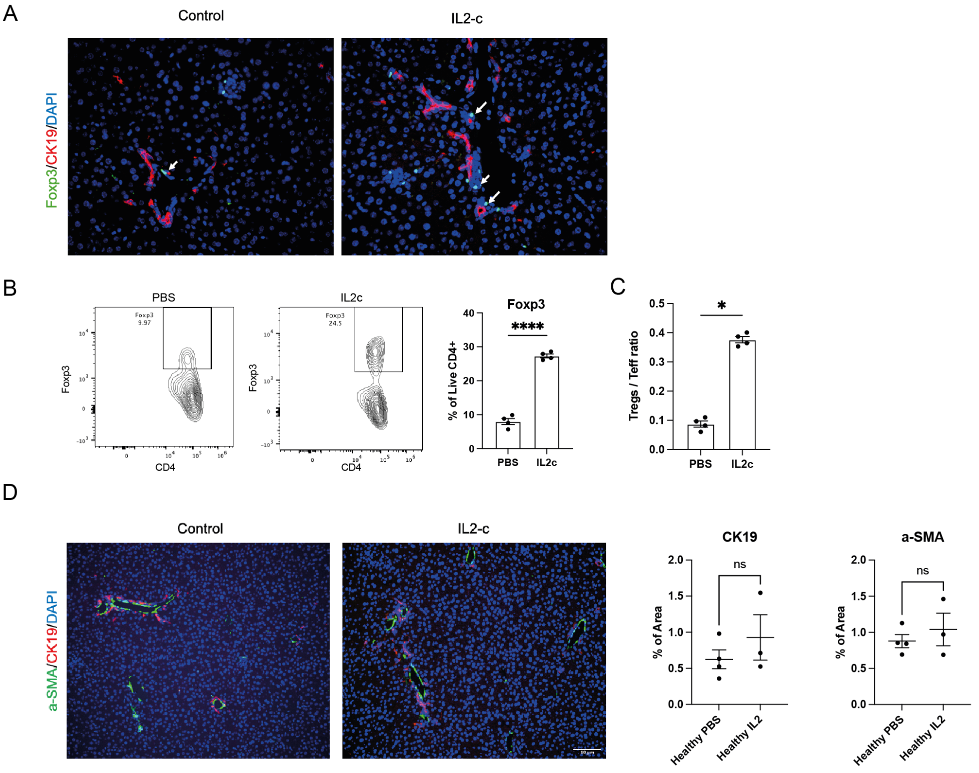


**Supplementary Figure 3. IL-2c injection expanded Foxp3+ Tregs in healthy mice.** A) Representative images of expanded Foxp3 Tregs (green)located near the CK19+ bile ducts (red). B ,Flow cytometry analysis and quantification of intrahepatic Foxp3 Tregs. C) Intrahepatic Tregs to T effector cells ratio after IL2c mediated Tregs expansion compared to control. D) Representative images and quantification of CK19+ ductular reaction (red), and aSMA+ activated fibroblasts (green).

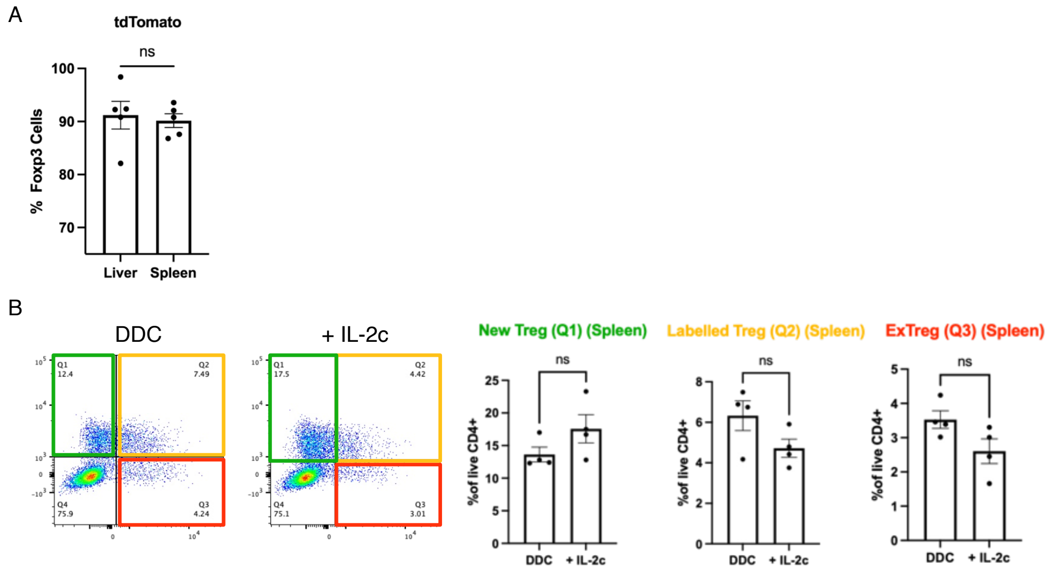


**Supplementary Figure 4. IL-2c injection did not increase the exTreg population in the spleen following DDC-induced liver injury.** A) Labelling efficiency of spleen and liver Tregs of the Foxp3$^{Cre}$Ai14 mice. B) Flow cytometry analysis of labelled Tregs populations in the spleen.

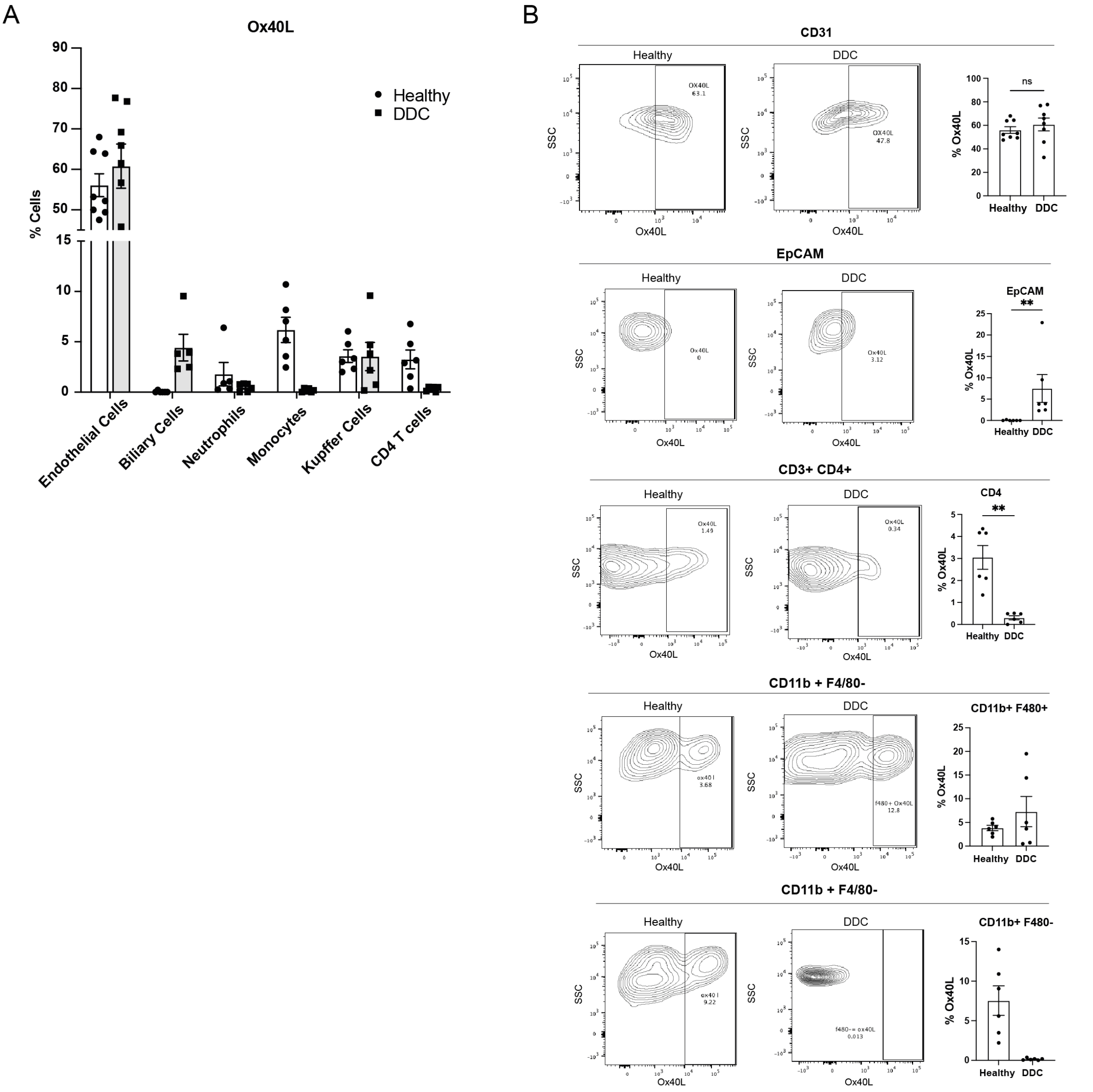


**Supplementary Figure 5. Ox40L expressing cells in the liver following DDC-induced injury.** A) Quantification of intrahepatic Ox40L expressing cells using flow cytometry analysis. B) Ox40L expression of CD31+ endothelial cells, EpCAM+ biliary cells, CD4+ T lymphocytes, CD11b$^{hi}$F4/80- monocytes, CD11b$^{lo}$ F4/80+ macrophages isolated from the liver.

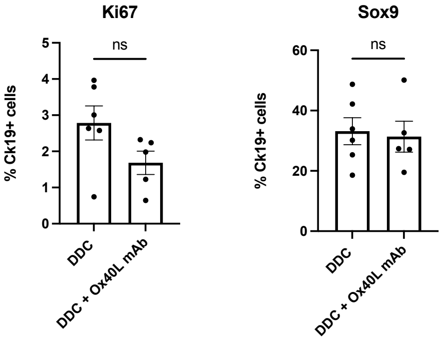


**Supplementary Figure 6. Ox40L administration does not affect cholangiocyte proliferation and the ability to form ductular reactions.** A) Quantification of Ki67 and Sox9 expressing Ck19+ biliary cells post Ox40L mAb administration following DDC-induced liver injury.

**Supplementary Table 1- Primary antibodies**

**Anti-mouse**

| Antibody used | Host | Cat Number | Dilution |
|---|---|---|---|
| CK19 | Rat | DSHB AB_2133570 | 1:400 |
| Foxp3 | Rabbit | 12653S | 1:200 |
| CD4 | Rabbit | Ab183685 | 1:300 |
| CD8 | Rabbit | Ab217344 | 1:300 |
| Sox9 | Rabbit | Ab5535 | 1:300 |
| Yap1 | Rabbit | 14074S | 1:200 |
| Gpnmb | Goat | AF2330 | 1:200 |
| RFP | Goat | 200-101-379 | 1:200 |
| OX40L | Rabbit | PA5-34516 | 1:200 |
| Iba-1 | Rat | Ab283346 | 1:200 |
| CD206 | Rabbit | NBP-1-90020 | 1:200 |
| T-bet | Rabbit | D6N8B | 1:200 |

**Anti-human**

| Antibody used | Host | Cat Number | Dilution |
|---|---|---|---|
| PanCK | Rabbit | Z0622 | 1:400 |
| CD4 | Goat | AF-379-NA | 1:500 |
| CD8 | Mouse | 66868-1-Ig | 1:100 |
| OX40L | Rabbit | PA5-80169 | 1:500 |

**Supplementary Table 2- Secondary antibodies**

| Antibody | Host | Cat Number | Dilution |
|---|---|---|---|
| Anti-rabbit 488 | Donkey | A21206 | 1:500 |
| Anti-rabbit 555 | Donkey | A31572 | 1:500 |
| Anti-rabbit 647 | Donkey | A31537 | 1:500 |
| Anti-rat 594 | Donkey | A21209 | 1:500 |
| Anti-rat 488 | Donkey | A21208 | 1:500 |
| Anti-mouse 555 | Goat | A21422 | 1:500 |
| Anti-goat 555 | Donkey | A21432 | 1:500 |
| Anti-goat 488 | Donkey | A11055 | 1:500 |

**Supplementary Table 3- Antibody for flow cytometry**

Antibody panel (all from BioLegend with dilution 1:200):

| Antibody | Colour | Clone |
|---|---|---|
| CD45 | Alexa Flour 700 | 30-F11 |
| CD3 | BV785 | 17A2 |
| CD4 | Pe-Cy5 | GK1.5 |
| CD8 | PerCP | 53-6.7 |
| Foxp3 | FITC | FJK-16s |
| T-bet | Pe-Cy7 | 4B10 |
| Ifng | BV510 | 505841 |
| OX40 | BV421 | 119411 |
| CD25 | BV605 | 102036 |
| Tnfa | BV711 | 563944 |
